# An industry perspective on whole genome-informed hybrid maize disease resistance characterization to improve breeding decisions

**DOI:** 10.64898/2026.07.30.741863

**Authors:** Jill C. Check, Frank Technow, Liviu Radu Totir

## Abstract

Characterizing hybrid maize disease resistance is a costly and labor-intensive effort in commercial breeding programs. Field trials are carefully inoculated and managed but remain error-prone due to spatial variability in disease pressure, microclimatic conditions and inter-rater variability. Quantitative ordinal disease rating scales are used to increase scoring speed at the expense of resolution, accuracy, and the ability to use conventional statistical methods. To improve traditional methods of disease resistance characterization, we propose to leverage readily available low-density SNP marker profiles to create genome-informed disease scores. Specifically, a whole genome ordered probit regression (WGOPR) model is used to deconstruct field-observed disease phenotypes into marker effects and reconstruct genome-informed disease scores. This approach is demonstrated in hybrid maize using data from *Exserohilum turcicum*-inoculated field trials across the central and northern U.S. and Canadian Corn Belt in 2024. Resulting Genomic Estimated Categorical Probabilities (GECPs) are compared to observed frequencies of disease scores to validate the methodology and evaluate the accuracy of regional hybrid maize disease resistance characterization. The benefit of a probabilistic output is demonstrated through two use cases: a comparison of hybrids with highly variable observed disease resistance scores at a single location, and a comparison of breeding selection schemes from a regional analysis. Because GECPs are the product of estimated marker effects, they better represent the expected behavior of a genotype independent of location-, rater- and plot-specific noise, and will therefore offer a step towards improving hybrid maize characterization and better informing breeding decisions.

## Introduction

With increasing global food demand (Tilman et al., 2011) and increasing threat of plant disease due to pathogen evolution, transmission via trade networks, and climate change (Ristaino et al., 2021), both yield and disease resistance are vital plant breeding objectives. While planting resistant genotypes is an important part of any integrated pest management program, the tradeoff between yield potential and disease resistance is well documented (Derbyshire et al., 2024; Ning et al., 2017; Smedegaardpetersen and Tolstrup, 1985). Therefore, plant breeders must make careful selections to balance both yield potential and yield stability under disease pressure, which requires confident characterization of both traits.

In typical commercial breeding pipelines, disease resistance characterization is performed by scoring the phenotypic reaction of a candidate in a network of inoculated field trials across a target population of environments (TPEs). To accomplish this, quantitative ordinal scales are used instead of near point severity estimates for disease assessment. Ordinal data is assigned into distinct categories with a natural ordering but often unequal or immeasurable differences between categories, which introduces uncertainty in both scoring resolution, accuracy and use of conventional statistical methods (Bock et al., 2024). An example of a quantitative ordinal scale is the Horsfall-Barratt (HB) scale, which is commonly used by pathologists and breeders to assess foliar disease severity (Guan et al. 2023; Strayer-Scherer et al. 2018; Weber and Jansky 2011). In the HB scale, percent severity is divided into 12 logarithmically scaled intervals symmetrical around 50% severity (Horsfall and Barrat 1945). Rank-based statistical analysis methods (Shah and Madden, 2004) or midpoint conversion (Bock et al., 2010) are used for hypothesis testing using uneven interval quantitative ordinal data. However, retaining the true value of the response variable is important for interpretation of disease resistance scores and selection of breeding candidates. Despite these known pitfalls, using quantitative ordinal disease rating scales is necessary given the large number of candidates being assessed in large scale breeding programs and the need to preserve consistent data structure with previous years of observations.

Decades of research have been dedicated to advancing statistical methods that accurately evaluate and characterize breeding candidates and improve selection decisions (Bernardo, 2021). Using the mixed linear models statistical framework to make within-environment spatial adjustments (Gilmour et al., 1997) and across-environment adjustments (Cooper et al., 2014) has improved the resolution of field phenotyping and has become the standard approach to evaluate and characterize the performance of breeding candidates in multi-environment trials. Building on the mathematical foundation of mixed linear model methodology, the seminal work by Meuwissen et al. (2001) marked a major leap forward, laying the foundation for genomic selection by integrating into mixed linear model methodology whole genome Single Nucleotide Polymorphism (SNP) information, generated through high-throughput genotyping technologies. This breakthrough allowed for the joint assessment of markers across the genome, supplanting the need for careful marker curation and QTL validation to assess the genetic value of a candidate for polygenic traits (de Koning, 2016). Recently, a large genomic selection validation study was conducted by Corteva Agriscience, demonstrating the value of whole-genome prediction methodologies in modern plant breeding (Jines et al., 2025). These models have broadly replaced the use of raw field observations in breeding decisions for traits represented by continuous scales (e.g., yield, moisture, test weight). However, there is presently no similar approach used for ordinal traits which violate the assumptions of conventional linear models, including disease resistance traits. Therefore, to best inform breeding decisions and progress both yield and disease resistance objectives, whole genome-informed statistical modeling methods are needed to improve the resolution of disease resistance characterization.

To address these limitations and improve existing methods of disease resistance characterization, a genomic-informed approach is employed. Whole genome ordered probit regression (WGOPR) is used to characterize hybrid maize performance using a network of *Exserohilum turcicum*-inoculated field trials for northern corn leaf blight (NCLB) resistance screening from the central and northern U.S. and Canadian Corn Belt. The approach is employed on two scales: for individual trial locations and for the full trial network. The credibility and accuracy of predictions from individual locations are compared and linked to the characteristics of their observed data. Potential selection schemes of breeding program candidates using the multi-environment trial data are explored. The benefits of probabilistic interpretation and potential candidate comparisons and selection schemes are demonstrated at both scales. Together, this framework provides a statistically rigorous approach for improving the accuracy and interpretability of ordinal disease resistance scores of candidates in commercial maize breeding programs.

## Materials and Methods

### Whole genome ordered probit regression (WGOPR)

Hybrid disease resistance is modeled using a Bayesian ordered probit regression framework. The following generalized linear model was fitted to the observed ordinal disease resistance scores, separately for each location:

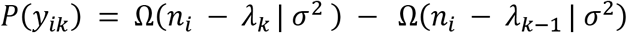

Where *P*(*y_ik_*) is the probability of observing disease score *k* in plot *i*, Ω denotes the Gaussian probability density function with mean *n_i_* and variance *σ*^2^and with *λ_k_* and *λ_k_*_−1_ being the thresholds demarcating category *k* in the latent probit space. The lowest and highest categories are bounded by *λ*_0_ = −∞ and *λ_K_* = ∞, respectively. The linear predictor *n_i_* was modeled as

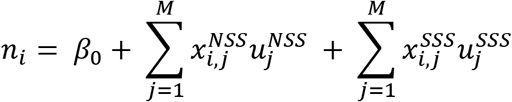

Where *β*_0_ is an intercept, 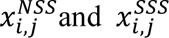 represent the centered genotype scores of marker *j* of the parents from each heterotic group (non-stiff stalk (NSS) and stiff stalk (SSS)) of the hybrid in plot *i*, and 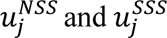 represent the corresponding marker effects. Prior distributions of marker effects correspond to those of the standard BayesB formulation (Meuwissen et al., 2001). The intercept was fixed to an arbitrary constant of 0 and *σ*^2^ to a value of 1 as both simply shift and scale the values of the thresholds *λ* but do not affect the fitted probabilities. The probability of the linear predictor *n_i_* that occupies the space between each pair of thresholds represents the probability of belonging to that category, as illustrated in **Figure 1**. The model was fitted using Markov Chain Monte Carlo (MCMC) sampling with 15000 iterations and 50% burn in.

**Figure 1:**
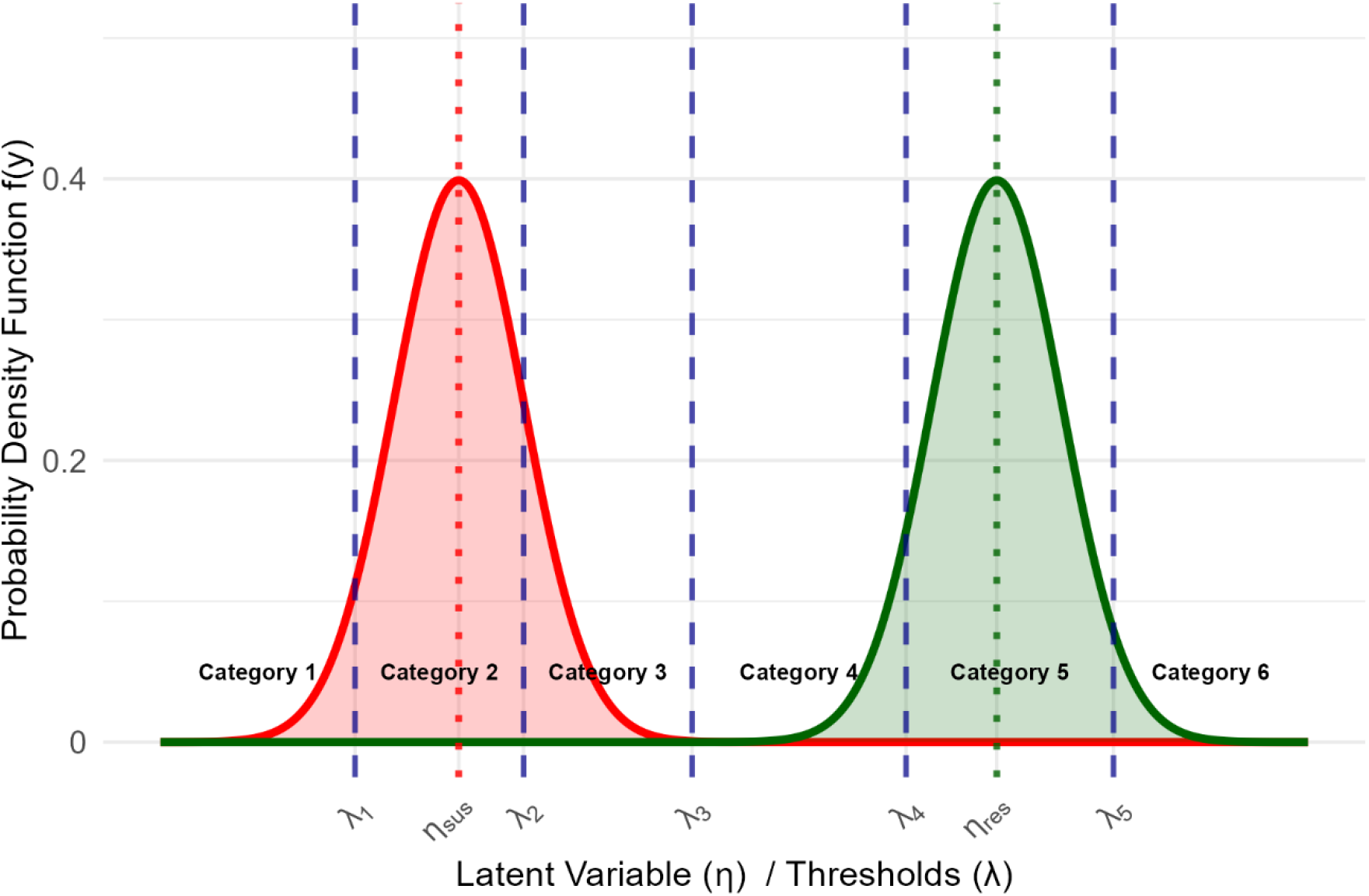
A visual representation of Genomic Estimated Categorical Probabilities (GECPs) for a susceptible maize hybrid (solid red line) with a latent disease score *n_sus_* (dotted red line) and a resistant maize hybrid (solid green line) with a latent disease score of *n_res_* (dotted green line). The GECPs of the two hybrids are represented by the probability density function for a normal distribution with mean of *n_i_* and *σ*^2^ of 1. Dashed blue lines demarcate the threshold values for the individual categories.

Furthermore, the model can be extended to characterize hybrid disease resistance scores from multi-environment trial data:

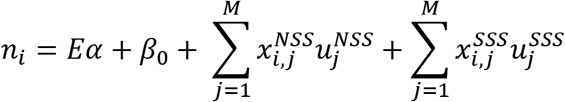

where *E* is the design matrix encoding location identifiers and *α* is a vector of location-specific effects. The location effects are assigned a Bayesian ridge regression prior distribution. Fitting location effects allows the model to capture systematic differences in disease pressure, which could be due to factors such as disease rating timing, inoculation quality, or incubation conditions, while maintaining the parent-specific genetic architecture of the hybrids. Technical details about fitting probit models for categorical data are described in Albert and Chibb, 1993.

Given these statistical models, the probability for all scored categories was calculated for each hybrid. The resulting probabilities of belonging to a given category *k* are referred herein as Genomic Estimated Categorical Probabilities (GECPs).

### Model assessment metrics

Credibility, calibration, and accuracy model assessment metrics of predicted GECPs were used to evaluate the ability of the WGOPR model to reconstruct the observed phenotypes using genomic data. Credibility was assessed using the Effective Number of Credible Categories (ENCC) per genotype, a novel statistic used to summarize the spread of predicted probabilities across ordinal categories into a single value using the complete probability vector. For example, a genotype with a probability vector (0.00, 0.00, 0.05, 0.19, 0.26, 0.27, 0.20, 0.03, 0.00) results in a greater ENCC than a genotype with a probability vector (0.00, 0.00, 0.00, 0.27, 0.50, 0.23, 0.00, 0.00, 0.00). ENCC is calculated as:

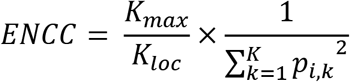

where K is the number of categories and *p_i_*_,*k*_ is the probability of individual i belonging to category k. Because locations differed in the number of categories scored, ENCC was scaled by the ratio of the total possible number of categories on the ordinal scale (*K_max_*) to the number of categories scored at the location (*K_loc_*). When all 9 categories were observed in the data, the scaling factor was 1. A lower ENCC value represents greater credibility in fewer categories, whereas a higher ENCC indicates more ambiguity in the classification across categories.

As is standard, model calibration was assessed by comparing the observed frequencies of categories to their predicted probabilities. A probabilistic model is considered calibrated, when the observed frequency of predicted outcomes matches their predicted probability. The observed frequency is calculated as the proportion of observations where the predicted category was observed to the total number of observations within a category and probability interval:

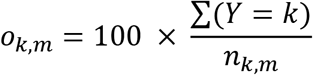

where *o_k_*_,*m*_ is the observed frequency for category k and probability interval bin m, *Y* = *k* is a logical operator indicating cases when observed and predicted categories match and *n_k_*_,*m*_ is the number of observations within category k and probability bin m. Four probability bins were used: (0, 15], (15, 30], (30, 45] and (45, 100]. The width of the bins were chosen such that the minimum number of observations within each was sufficient for accurately calculating the observed frequencies (50). For example, an observed frequency of 10% for category 3 and probability bin (0, 15] would indicate that 10% of individuals with a predicted probability between 0 and 15% for category 3 were observed as 3. Expected Calibration Error (ECE) (Nixon et al. 2019) is used to summarize the difference between the observed frequency and predicted probabilities across categories and probability intervals. ECE is calculated as:

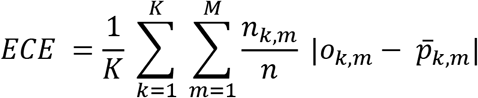

where M is the total number of probability bins, n is the total number of observations and *p_k_*_,*m*_ is the average probability of observations in the probability bin m and category k. Differences between observed frequencies (*o_k_*_,*m*_) and average predicted probabilities (*p_k_*_,*m*_) are weighted by the number of observations per category and probability bin. Division by K scales the values between 0 and 1, where a lower ECE indicates better prediction.

Model accuracy was assessed using the Ranked Probability Score (RPS) of each genotype. Like ECE, RPS measures the difference between observed scores and GECPs, but RPS penalizes predictions based on the distance between the observed category and highly predicted categorical probabilities (Epstein 1969). RPS also offers a measure of model accuracy that is insensitive to the binning of probabilities, unlike ECE. RPS represents the difference between the observed category and the cumulative predicted probabilities as:

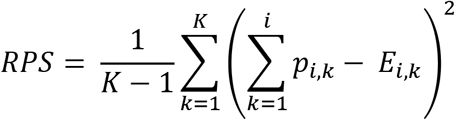

where *p_i_*_,*k*_ represents the probability of individual i belonging to category k and *E_i_*_,*k*_ is an indicator variable, where *E_i_*_,*k*_ = 1 when the observed category is less than or equal to k for individual i, and *E_i_*_,*k*_ = 0 otherwise. The normalization factor ensures RPS ranges from 0 (perfect prediction) to 1 (maximum possible error, e.g., predicting category 1 when category 9 was observed). A lower RPS indicates closer concordance between predicted probabilities and observed scores.

### Demonstration data set description

The proposed technical solution is demonstrated using an example data set comprised of seven hybrid maize field trials grown across the northern and central U.S. and Canadian Corn Belt in 2024 (**Table 1**). Relative maturity groups from 85 to 110 are represented. Field trials were inoculated with *E. turcicum* to characterize the disease resistance of pre-commercial maize hybrids to northern corn leaf blight (NCLB). Disease severity was assessed on a whole-plot basis. Disease severity was scored on a quantitative, uneven interval ordinal scale from 1 to 9, where 1 = >80% severity, 2 = 66-80% severity, 3 = 51-65% severity, 4 = 36-50% severity, 5 = 26-35% severity, 6 = 16-25% severity, 7 = 6-15% severity, 8 = 2-5% severity and 9 = < 2% severity. A set of well-characterized hybrid maize checks are used to determine the timing of disease resistance scoring. When checks reach their expected disease resistance scores, all maize hybrids are scored. This method is used in an effort to control the effect of rating timing on disease resistance scores. Field trial location characteristics and hybrid population summary statistics are detailed in **Table 1**.

**Table 1:** Description of *Exserohilum turcicum*-inoculated hybrid maize field trial locations used in this study.

| Location code | Municipality | Planting date | Irrigated | Number of plots | Number of hybrids | Mean observations per hybrid | Unique SSS parents | Unique NSS parents | Mean crosses per parent |
| --- | --- | --- | --- | --- | --- | --- | --- | --- | --- |
| IA1 | Polk, IA | 5/29/2024 | Yes | 1405 | 697 | 2.02 | 283 | 256 | 5.23 |
| IA2 | Kossuth, IA | 5/15/2024 | No | 2849 | 2102 | 1.36 | 430 | 435 | 6.59 |
| MI1 | Gratiot, MI | 5/25/2024 | Yes | 3490 | 2004 | 1.74 | 439 | 374 | 8.64 |
| PA1 | Lancaster, PA | 5/19/2024 | Yes | 4114 | 3273 | 1.26 | 355 | 351 | 11.66 |
| PA2 | Lancaster, PA | 5/22/2024 | Yes | 4135 | 1831 | 2.26 | 537 | 459 | 8.36 |
| QC1 | Soulanges, QC | 5/22/2024 | No | 1754 | 650 | 2.70 | 249 | 261 | 6.88 |
| WI1 | Eau Claire, WI | 5/18/2024 | No | 3595 | 1863 | 1.93 | 404 | 394 | 9.01 |

The demonstration data was analyzed in two ways: on a single-field basis and on a regional basis. In both cases, the same criteria were used to assess GECP predictions (ENCC, ECE and RPS). The quality of GECP predictions from individual locations were compared and contrasted. If fitted marker effects cannot reconstruct the field-observed phenotypes at a given field location, it is understood that associations between phenotypes and genomic data are hidden by external noise, and that data from these field locations may be unreliable. Differences in the reconstruction of field-observed phenotypes between field locations were related to the characteristics of their observed disease scores. Data from all field locations were used to create regional disease resistance scores for all maize hybrids, which better reflect a commercial breeding program’s approach to disease characterization in comparison to analysis of single-location data.

All inbred parents of the hybrids present in the demonstration dataset were genotyped with a whole genome prediction SNP panel that ensured appropriate whole genome coverage given the linkage and linkage disequilibrium structure of the germplasm in question.

### Code and data availability

All computations were performed in the R statistical computing environment (R Core Team 2025). The probit model was fitted using the BGLR R package (Perez and de los Campos, 2014). As the NCLB disease screening network is a part of Corteva Agriscience’s commercial breeding program, the genetic and pedigree data constitute trade secrets and cannot be made available.

## Results

### Single-location analyses

The Genomic Estimated Categorical Probabilities (GECPs) generally aligned with their observed disease scores at each location. This alignment is visible as a progressive shift of the probability mass toward higher ordinal categories as observed disease scores increase from susceptible (low ordinal categories) to resistant (high ordinal categories) (**Fig. 2**). However, the seven locations differed greatly in their distribution of field-observed disease scores. The WI1 location more frequently scored susceptible phenotypes, IA2 more frequently scored resistant phenotypes, and IA1 more heavily scored intermediate phenotypes (central ordinal categories). Locations MI1, PA1, PA2 and QC1 had the most complete representation of observed ordinal categories with at least 8 of 9 ordinal categories observed (**Table 2**). The limited phenotype diversity in observed scores at some locations hindered the genomic model’s ability to fit meaningful marker effects. This is illustrated by the concentration of highly predicted probabilities at IA1 and IA2 around a few central categories, indicating shrinkage of GECPs in the absence of meaningful genomic relationships. However, at WI1, a lack of diversity in observed disease scores was reflected as low probability predictions across many categories. The low probability predictions across observed categories at WI1 are also reflected in the distribution of the Effective Number of Credible Categories (ENCC) (**Fig. 3**). WI1 had the greatest average ENCC across all locations (4.29; **Table 2**), indicating low credibility in GECPs and ambiguous classification across categories. WI1 was followed by IA2 and IA1, with mean ENCCs of 3.85 and 3.29, respectively. PA2 demonstrated the greatest credibility among locations, with a mean ENCC of 2.60.

**Figure 2:**
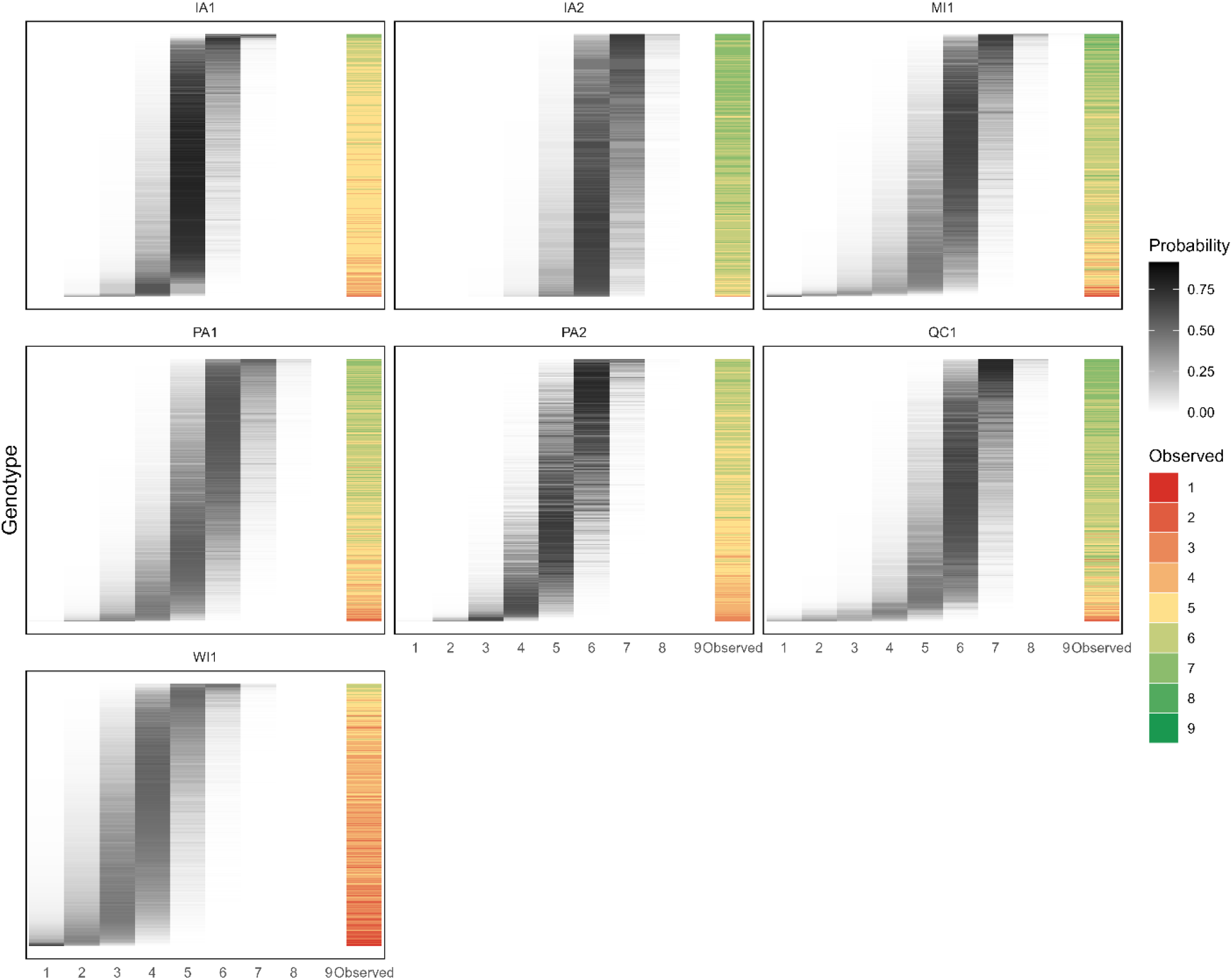
The predicted Genomic Estimated Categorical Probability (gray scale) distributions and the observed disease scores (red-yellow-green gradient scale) for each genotype within a location. Homogeneity in predicted probability distributions across genotypes indicates a lack of ability to reconstruct the observed disease phenotype using the genomic data. A visible “stair-step” pattern from low to high ordinal categories is indicative of the genomic model’s ability to distinguish between disease susceptibility categories. Poor model calibration is reflected in locations with a high degree of homogeneity in predicted probability distributions.

**Figure 3:**
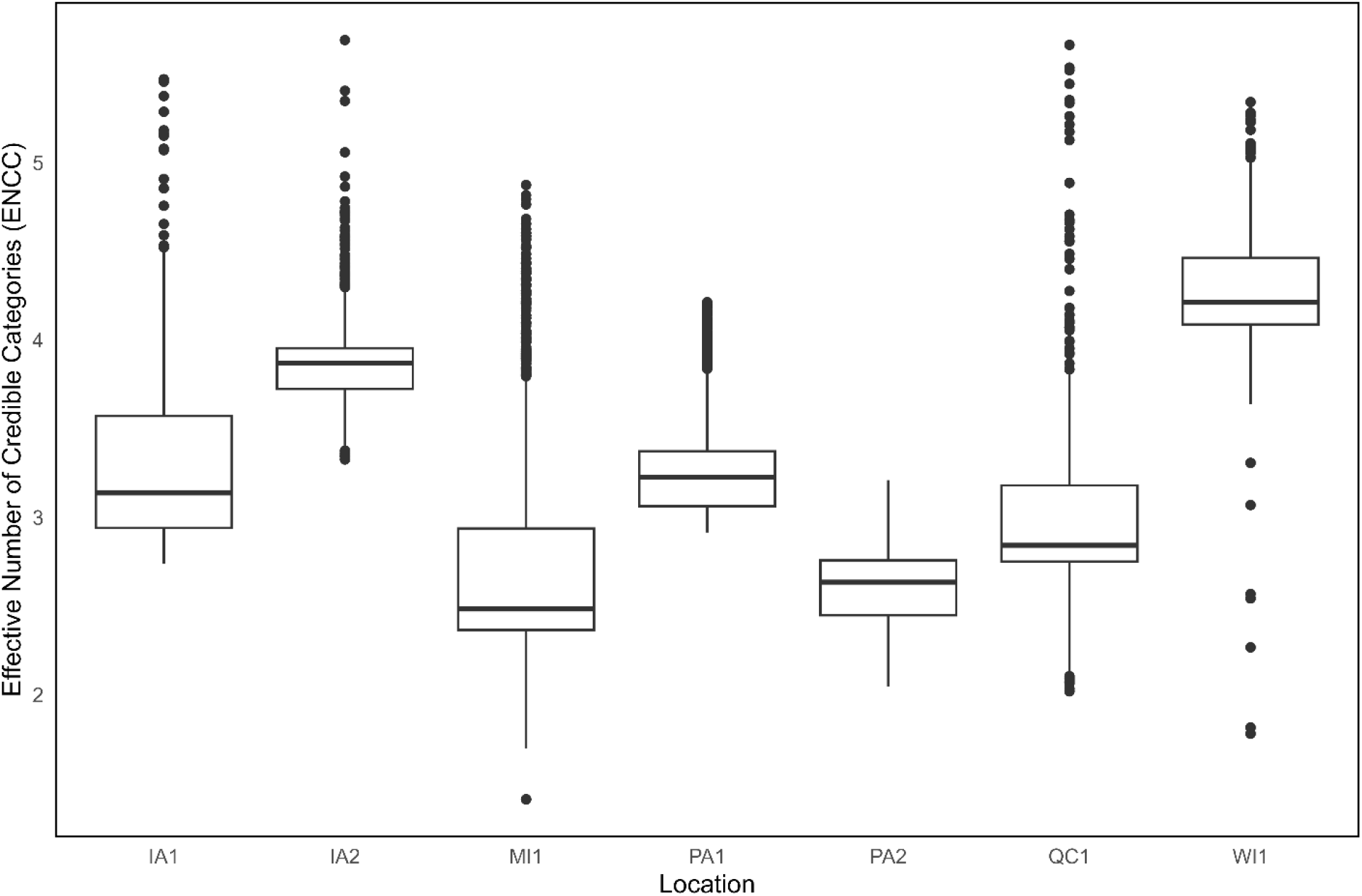
The distributions of Effective Number of Credible Categories (ENCC) for each location. ENCC is calculated for each genotype at each location. ENCC is a measure of prediction uncertainty and penalizes probability predictions based on their spread across categories. ENCC is scaled by the ratio of the number of total possible categories (9) to the number of observed categories per location. A lower location average ENCC indicates greater confidence in fewer categorical probabilities.

**Table 2:**
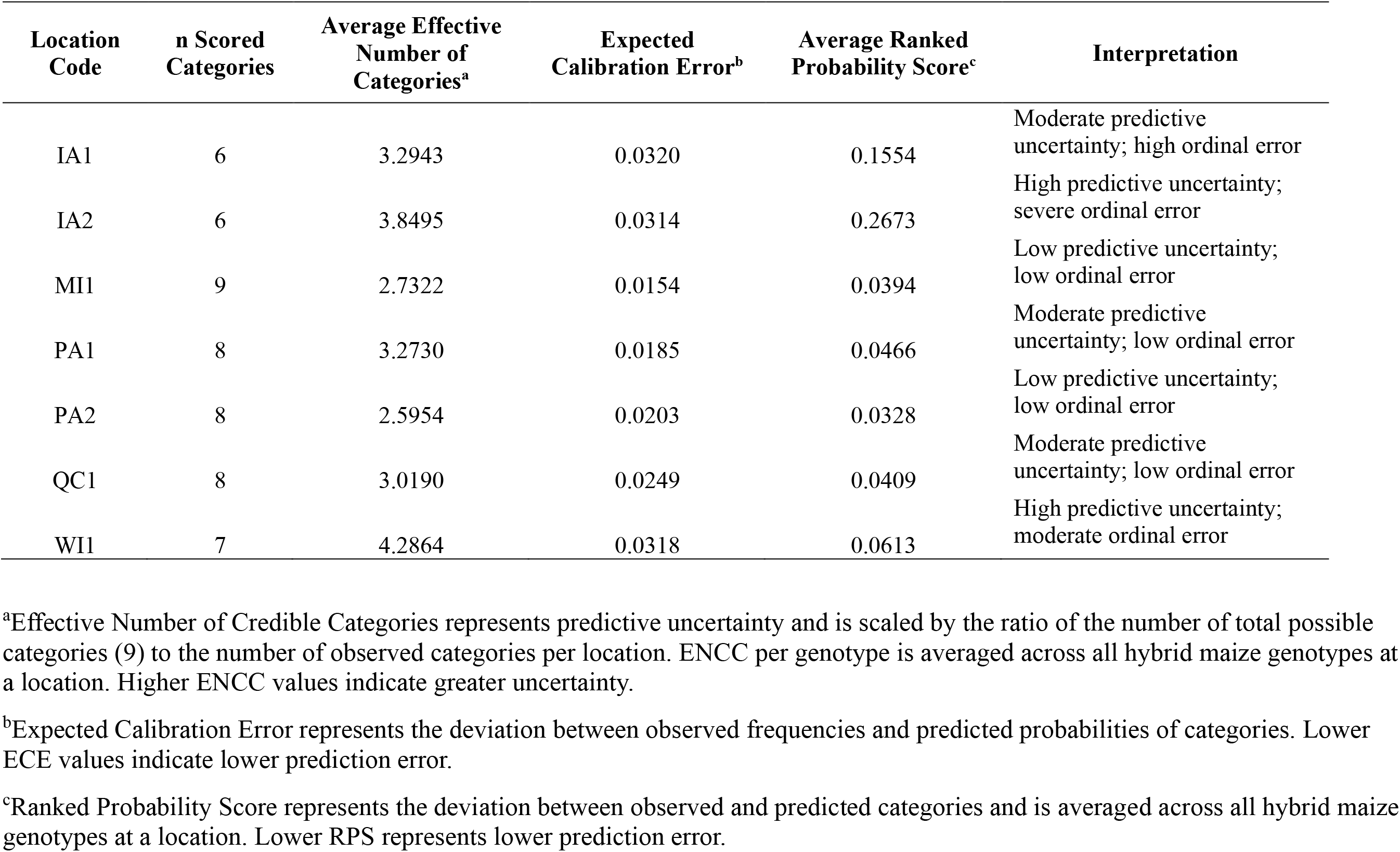
Single location observed categories, WGR model probability prediction statistics, and interpretations of location data characteristics.

The calibration of the Whole Genome Ordered Probit Regression (WGOPR) was assessed for each location (**Fig. 4**). GECPs are considered well-calibrated when the difference between the observed frequencies and average predicted probability are minimized for each category and probability interval bin. At locations PA1 and PA2, the observed frequences and average predicted probabilities deviated minimally except for lower categories where the model tended to underpredict probabilities. In contrast, IA1 and IA2 displayed substantial calibration errors. At IA1, the model concentrated probability predictions around category 5, where 1219 of 1405 genotypes predicted probabilities were between 45 and 100% probability. Similarly, probability predictions were concentrated around category 6 at IA2, where 2209 of 2849 genotypes predicted probabilities were between 45 and 100%. This concentration around a few highly predicted categories led to systematic underestimation in other categories and probability intervals. For example, at IA1, there was a 13.3 and 16.6% underestimation for the 15-30 and 30-45% probability intervals, respectively, for category 5. Calibration of GECPs was summarized by the Expected Calibration Error (ECE), which weighs the absolute differences between the observed frequency and the average predicted probability by the number of observations within each category and probability bin. ECE was greatest at IA1 (0.032), WI1 (0.32) and IA2 (0.31) (**Table 2**).

**Figure 4:**
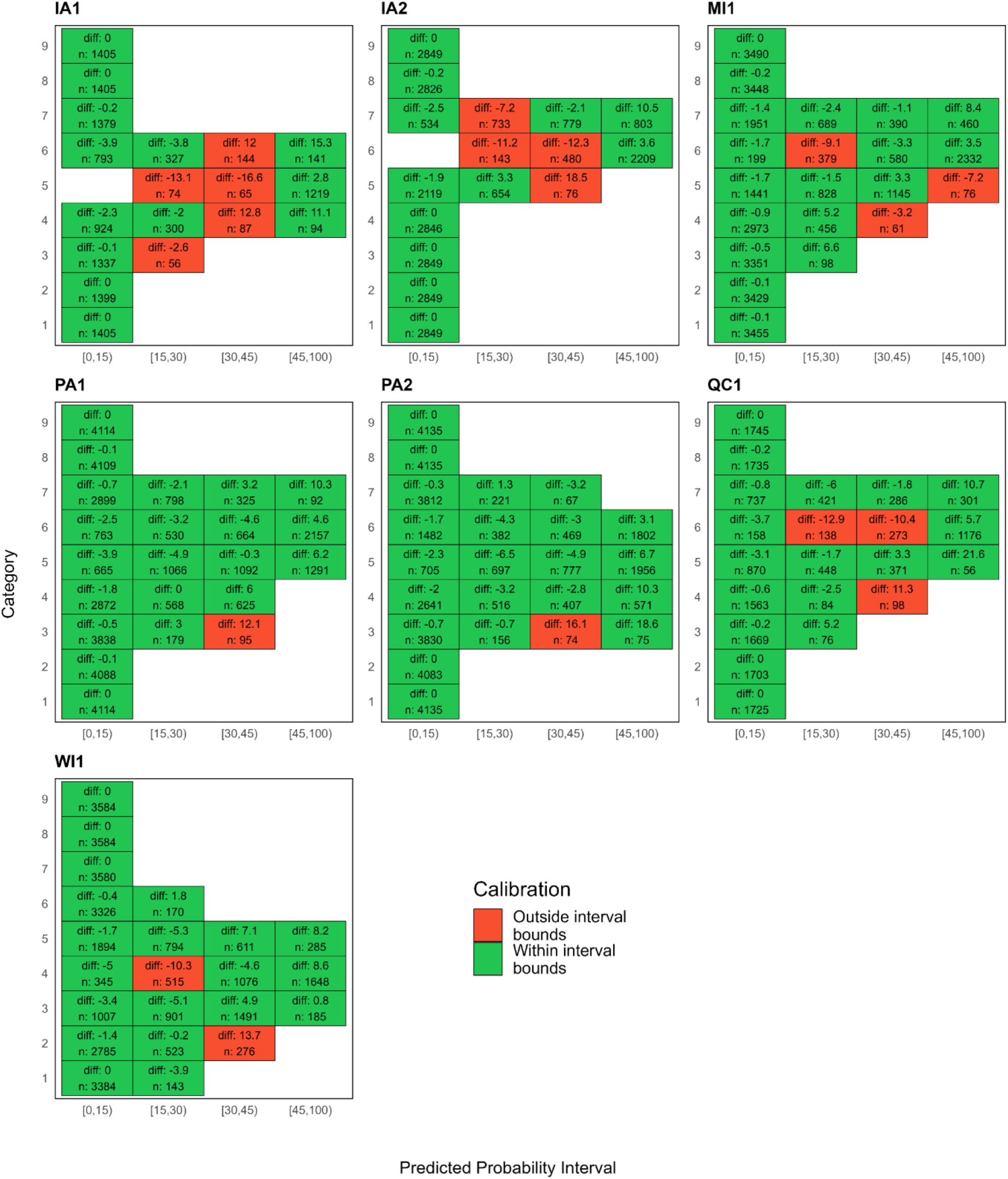
Single-location Genomic Estimated Categorical Probability calibration tables. Predicted probability intervals are compared to observed frequencies for each category with at least 50 observations. Predictions are considered calibrated when difference between the observed frequencies and the average predicted probability for each category and probability bin are minimized. Cells are labeled with the difference between the observed frequency and the average predicted probability (diff) and the number of observations (n) for each category and probability bin. Cells are colored based on whether the observed frequency falls outside or within the probability bin interval bounds.

Like ECE, Ranked Probability Scores (RPS) measures the deviation of predicted probabilities from observed categories. However, RPS penalizes based on the distance along the ordinal scale between the observed category and highly predicted probabilities, and is insensitive to bin bounds, unlike ECE. The mean and distribution of RPS were similar across all locations (0.03 to 0.06), except for IA1 and IA2, which had mean RPSs of 0.16 and 0.27, respectively, demonstrating substantial error in ordinal category predictions at these two locations (**Fig. 5**; **Table 2**). Despite having an ECE ranking like IA1 and IA2, WI1 had a substantially lower RPS (0.06). Together with the high ENCC estimate for WI1 (4.29), this indicates the model successfully employed the strengths of the Bayesian framework by spreading its uncertainty across a range of neighboring categories to avoid confident misses despite high environmental noise. This behavior contrasts with IA1 and IA2, which exhibited high ECE, high RPS but lower ENCC values, indicating a systemic ordinal mismatch where the model’s predictions were confidently distant from the observed disease phenotypes.

**Figure 5:**
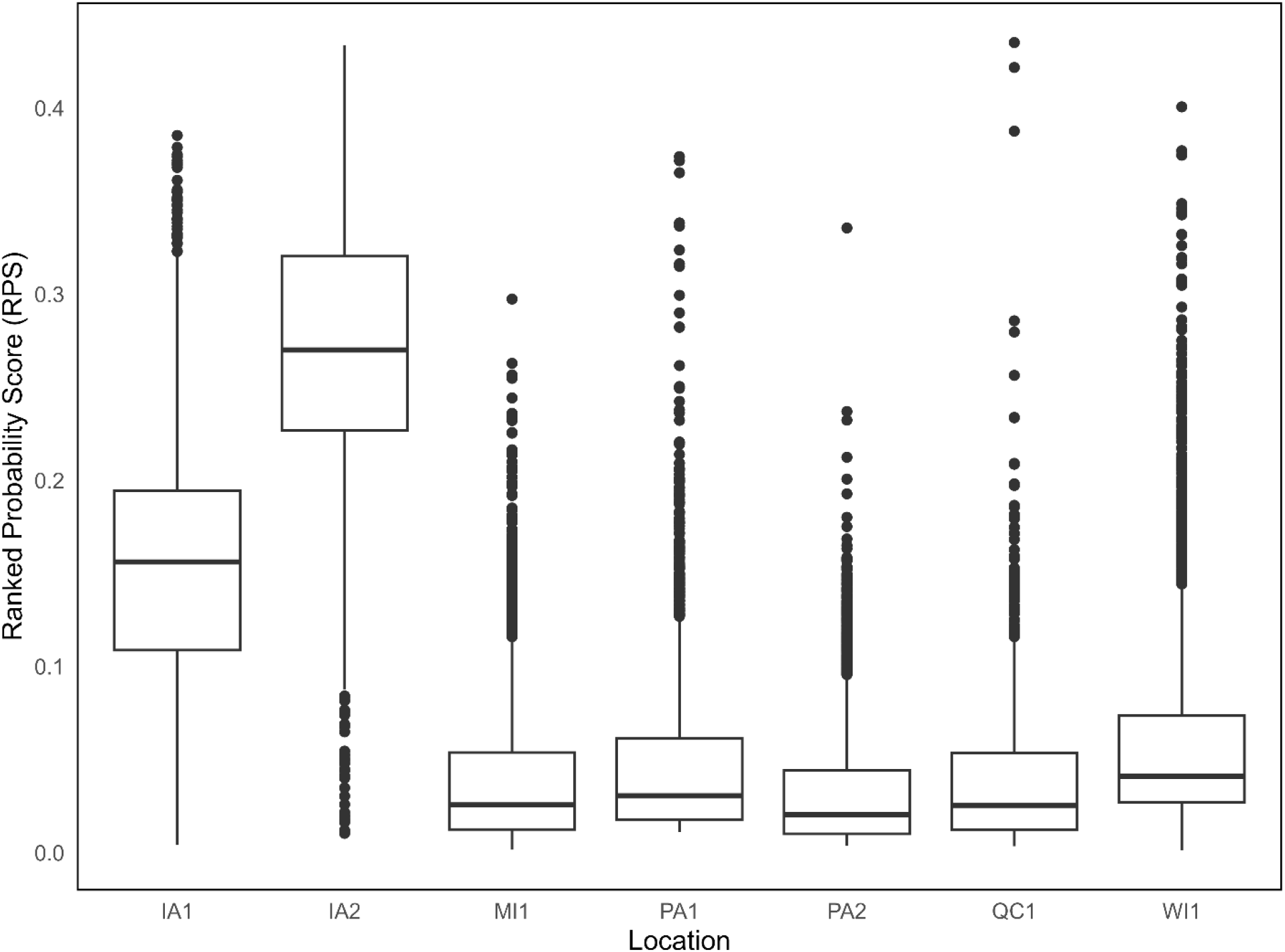
The distributions of Ranked Probability Scores (RPSs) for each location. RPS is calculated for each genotype at each location. RPS is a measure of prediction calibration error and penalizes predictions based on the distance between the observed and predicted ordinal categories (i.e., observed category 1 versus high predicted probability in category 9 is penalized more heavily than observed category 3 versus high predicted probability in category 4). A lower location mean RPS indicates closer concordance between predicted and observed scores.

### Regional analysis

To determine regional GECPs of all hybrid maize lines planted in the network of field trials in the demonstration data set, data from all locations were used in the multi-environment model. The use of multi-environmental trial data best represents an approach that would be used for disease characterization in a commercial breeding program. Despite localized systematic issues identified in the single-location analysis, the regional model was well-calibrated (**Fig. 6**). The regional model had a low ECE (0.18) and mean RPS (0.07) relative to the single-location models (**Fig. 7**). The regional model assigned high credibility to fewer categories, with a mean ENCC of 2.74, demonstrating that the predicted disease outcomes for most genotypes can be constrained to three or fewer ordinal categories. The credibility and accuracy of GECP predictions from the regional model underscore the robustness of the WGOPR approach, as the inclusion of multi-environment data mitigated the impact of low-quality locations.

**Figure 6:**
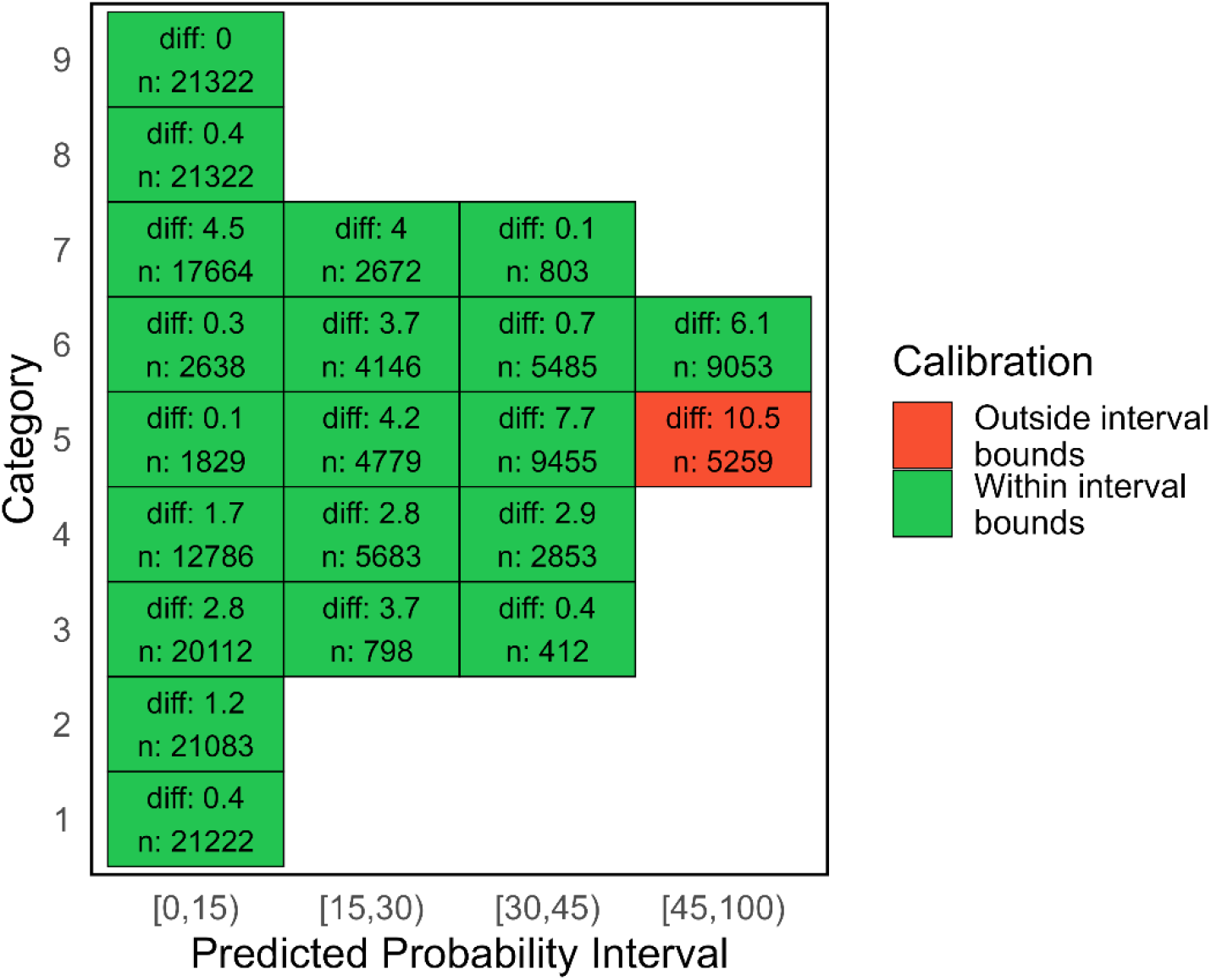
Genomic Estimated Categorical Probability calibration table for the regional model for hybrid maize disease resistance characterization. Predicted probability intervals are compared to the observed frequencies for each category with at least 250 observations. Predictions are considered calibrated when differences between observed frequencies and the average predicted probability for each category and probability bin are minimized. Cells are labeled with the difference between the observed frequency and the average predicted probability (diff) and the number of observations (n) for each category and probability bin. Cells are colored based on whether the observed frequency falls outside or within the probability bin interval bounds.

**Figure 7:**
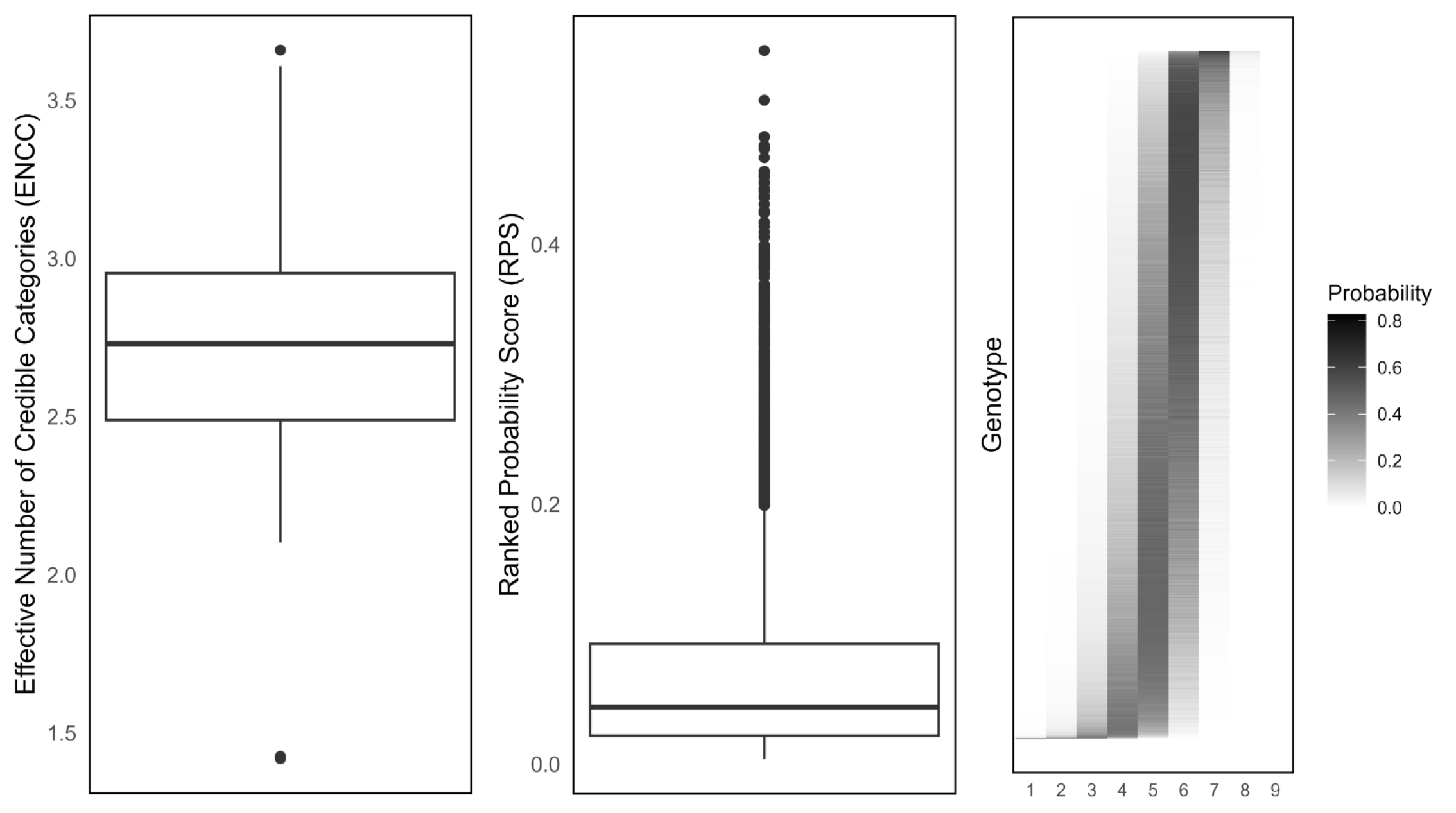
The distribution of Effective Number of Credible Categories (ENCC) for all genotypes included in the regional model (left). ENCC represents the spread of the probability distribution for each genotype across all categories on the ordinal scale. A lower ENCC represents high confidence (probability) across fewer categories and a higher ENCC represents low confidence (probability) across more categories. Lower ENCCs are more informative in disease susceptibility risk characterization. The distribution of Ranked Probability Scores (RPSs) for all genotypes included in the regional model (center). RPS is a measure of prediction calibration error, but penalizes predictions based on the distance between the observed and predicted ordinal categories. A lower location mean RPS indicates closer concordance between predicted and observed scores. Probability gradient for all genotypes in the regional model (right).

### Demonstrated use cases

A primary benefit of using WGOPR for characterizing disease resistance is the ability to interpret disease scores based on their categorical probabilities instead of single point estimates of ordinal categories for disease resistance scores. The advantage of this is that the categories have literal meanings, and the likelihood of observing certain levels of disease severity is informed by the GECPs. For example, category 6 represents between 16 and 25% NCLB foliar disease severity. Therefore, a hybrid with a 45% probability of belonging to category 6 can be expected to develop this level of severity under similar disease conditions in 45% of cases. Four hybrids from WI1 were selected to show the advantage of probabilistic interpretation using the demonstration data (**Fig. 8**). Each hybrid chosen had a high degree of variability in observed disease scores, with at least 4 different scores assigned to the same hybrid maize line between in-field replicates. The variability of observed disease scores highlights the complications of assigning single disease scores to hybrid maize lines. With the proposed approach, the underlying distribution of observed scores can be captured by the GECPs. For example, susceptible hybrid A was most often scored as category 2, but was also scored as a 1, 3 and 4. This is reflected in the probability distribution, where hybrid A had the greatest probability of belonging to category 2 (41%), followed by category 3 (32%), category 1 (21%) and category 4 (6%). Similarly, moderately resistant hybrid B was most often scored as category 5 and has a majority probability of belonging to category 5 (51%), followed by category 4 (28%), category 6 (18%) and category 3 (2%). Hybrid C and Hybrid D both represent moderately susceptible hybrids with very similar distributions of observed disease scores. However, the probability distributions allow for a richer inference in their predicted performances, where hybrid C has a 14% greater probability of belonging to categories 4 and above than hybrid D. This substantially higher risk associated with hybrid C compared to hybrid D cannot be distinguished when using point estimates of disease resistance scores.

**Figure 8:**
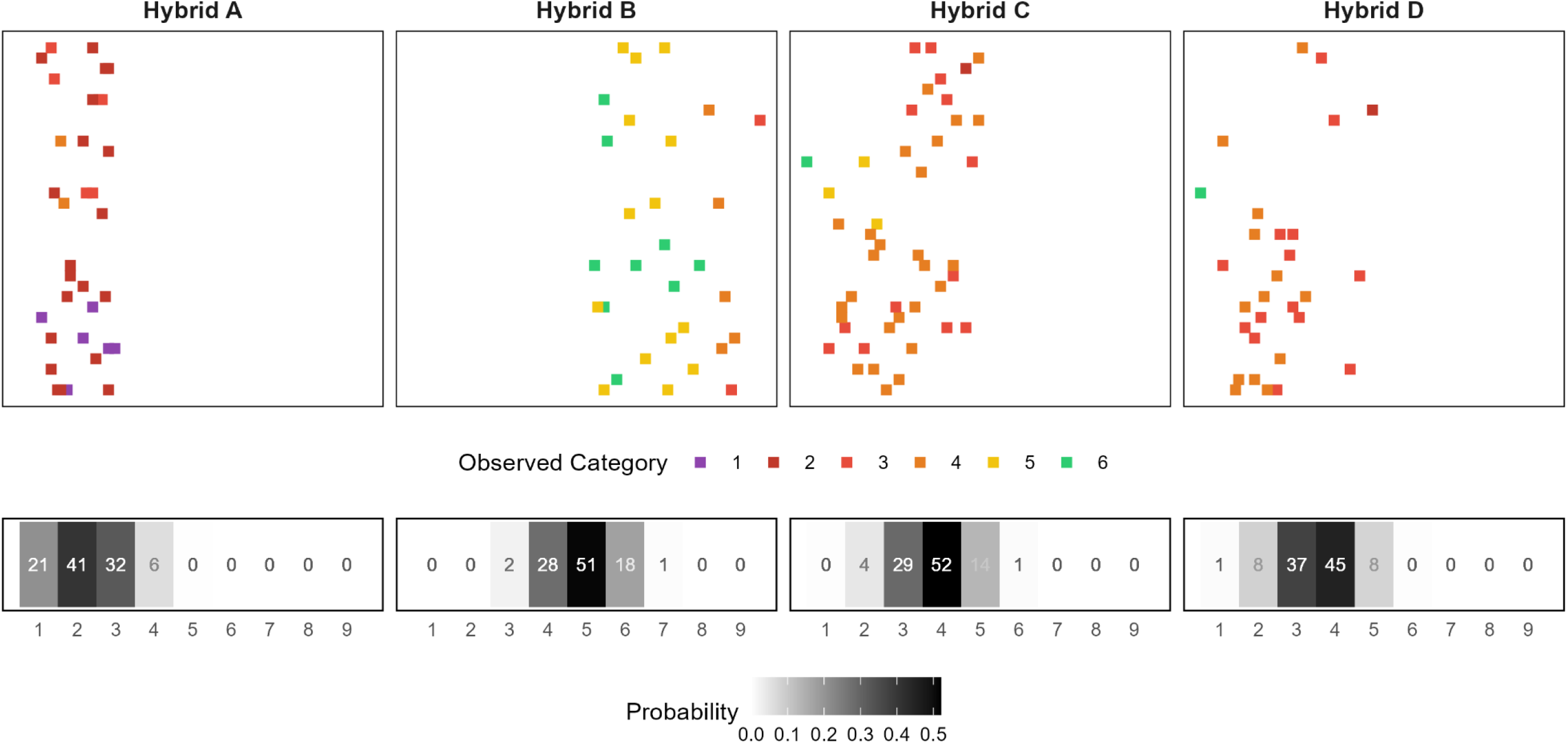
Four hybrid maize genotypes planted at WI1 with highly variable observed disease scores across in-field replicates (top) and their corresponding probability gradients across the ordinal scale (bottom). Hybrid A and Hybrid B represent susceptible and resistant genotypes, respectively. Hybrid C and Hybrid D represent moderately resistant hybrids with relatively different probability profiles. Hybrid C and D have the same mode (4) and similar ranges in observed disease scores, but the probabilistic approach allows for greater inference for the comparison of genotypes.

A common selection strategy is to discard individuals that are likely to be highly susceptible. The regional disease characterization results were used to demonstrate potential outcomes of different selection strategies for the demonstration data set (**Fig. 9**). The cumulative probabilities of categories 1 to 4 were used to represent the likelihood of observing a susceptible reaction. Three thresholds were defined to bin hybrid maize lines into low (0-25% probability), moderate (25-50%) and high (>50%) risk of being highly susceptible. A more conservative selection scheme may remove any candidate with 25% or greater probability of being highly susceptible. Employing this risk threshold would result in removing 1893 hybrid maize lines from the candidate pool. A less conservative selection scheme may only remove candidates with 50% or greater probability, removing only 523 candidates. By using probabilistic thresholds on GECPs, a maximum permissible risk can be used to determine breeding candidate removal, shifting selection strategies to focusing on the “safest” hybrids rather than just those with best performing averages.

**Figure 9:**
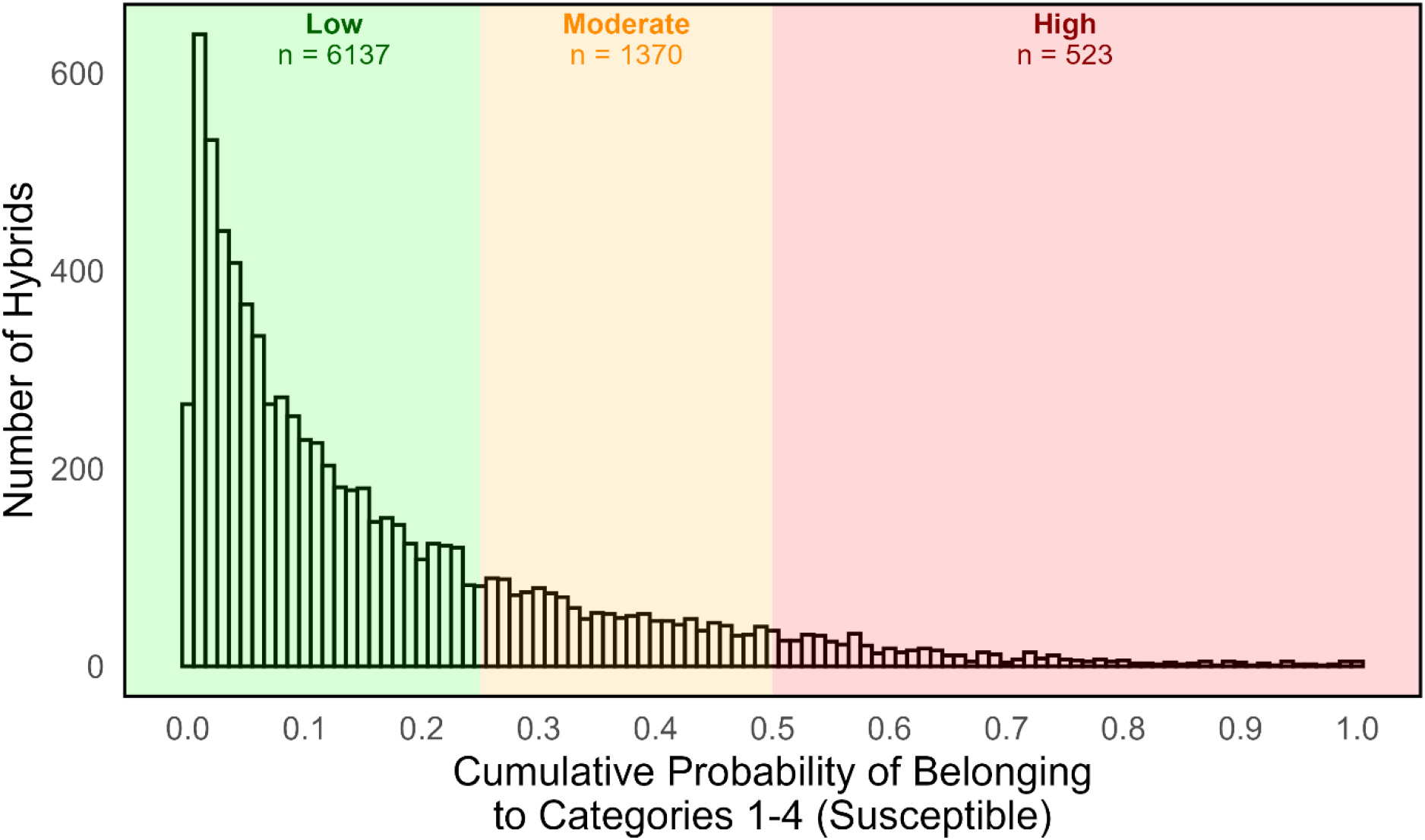
An example selection scheme for hybrid maize breeding advancements informed by a probabilistic approach. Hybrids are ranked in descending order based on the cumulative Genomic Estimated Categorical Probability for categories 1, 2, 3 and 4. Hybrids with a less than 25%, a 25-50%, or over 50% probability of being susceptible are considered low, moderate, or high risk for disease development under similar conditions, respectively.

## Discussion

High-quality disease resistance characterization is dependent on the availability of high-quality phenotype data. Spatial variability, inter-rater variability and inconsistency in inoculum viability and/or application are among many factors that may obscure the pure genetic response of an individual to pathogenic challenge in field trials. Despite best efforts by trial managers during field preparation, inoculation, management and data collection, these factors are difficult to control. Therefore, in an effort to best characterize hybrid maize disease resistance, we have demonstrated a genomic approach that enables a probabilistic interpretation of disease resistance and presented methods to evaluate the credibility and accuracy of GECPs as appropriate for the ordinal data. We used a network of NCLB disease resistance screening field trials as demonstration data to describe the methodology and dissect predictions from a range of locations. The quality of GECPs from individual locations was related to characteristics of the location’s observed data to highlight potential pitfalls in the collection of disease resistance phenotype data. Additionally, we demonstrated the robustness of the approach by leveraging the complete field trial network into a multi-environment model that produced GECPs equivalently credible and accurate to the best single-location models.

Overall, model performance varied significantly across the seven field trial locations. A core group of high-quality sites, characterized by low ECE and RPS were identified (MI1, PA1, PA2, QC1). At these locations, the relationship between disease resistance phenotypes and genomic data was strong enough to satisfactorily reconstruct the observed data. This core group shared similarities in their observed data that distinguished them from the other locations. At the high-quality sites, at least 8 of 9 ordinal categories were represented in the observed data, and the distribution of the observed scores was relatively balanced. For proper estimation of marker effects, contrasting disease phenotypes need to be observed in the data. Disease intensity and distribution are directly influenced by inoculation, incubation and trial management, highlighting the importance of pathology expertise in commercial breeding programs.

In commercial breeding programs, selection often focuses on removing highly susceptible individuals rather than advancing highly resistant individuals. This selection strategy is favored because genes governing defense networks have pleiotropic effects on plant growth and development, leading to tradeoffs between disease resistance and yield (Brown and Rant, 2013). Discarding only highly susceptible individuals leaves room to select for other traits of interest, especially yield. To effectively carry out this selection strategy, disease resistance scores need to be directly interpretable on the original ordinal scale in which they are collected. This requirement highlights the key limitation of using standard linear models to evaluate ordinal traits, which assume continuity and equal distance between categories. The WGOPR framework preserves the original meaning of the quantitative ordinal scale used for scoring disease resistance in the resulting GECPs while also allowing for a probabilistic interpretation. The probabilistic interpretation offers greater inference compared to single point estimates of disease resistance scores. Characterizing each candidate by categorical probabilities enables selection schemes based on cumulative probability thresholds (i.e., discard any individual with a 75% or greater probability of being highly susceptible) or between-category comparisons of individuals (i.e., what is the likelihood of belonging to category 1 versus 2?). Additionally, the original scale is familiar across stakeholder groups and retaining the ability to interpret disease resistance scores on this scale is important for communicating across expertise groups, such as plant breeders, pathologists, agronomists and salespeople. Most importantly, this ordinal scale is used to communicate disease resistance of commercial seed products to farmers. By maintaining the true meaning of the ordinal scale through product development, characterization and commercialization, a transparent understanding of disease resistance is conserved from end-to-end of the pipeline.

While predictive breeding, the assessment of candidates without direct observation of their phenotypes, was not the focus of this demonstration, the approach described potentially offers great advantages during data curation for prediction sets. As argued by Bernardo (2021), the success of predictive breeding will rely on the availability of high-quality phenotype data. The quality of predictions from genomic models are conditional on the ability to relate observed phenotypes to genetic factors. Thus, data quality should be defined by how effectively the measured phenotype captures genetic potential, which can be measured using the WGOPR approach used in this study. In the absence of genomic data, parental inbred parent main effects could be fit in place of marker effects. However, unlike the marker-effect approach used here, a pedigree-based approach would require a higher degree of replication of inbred parents and hybrids and be expected to have poorer accuracy (Crossa et al. 2010).

The demonstration of the WGOPR approach described herein was limited to a multi-environment inoculated trial characterizing NCLB disease resistance, but it is expected that the model utility and complexity will depend on characteristics of the plant-pathogen system. The genetic architecture of disease resistance will determine the model’s ability to estimate marker effects and thereby the success of reconstructing field-observed phenotypes. For example, maize resistance to NCLB is controlled by both qualitative resistances, conferred by *Ht* genes, and quantitative resistances, with little evidence of significant epistatic interactions (Chen et al. 2015; Chung et al. 2011; Welz and Geiger 1999). In plant-pathogen systems greatly influenced by epistatic interactions of quantitative resistance sources, the model may require additional complexity to satisfactorily reconstruct field-observed phenotypes based on additive gene effects only. Extensions of the original threshold model have been explored by Montesinos-Lopez et al. 2015, including additive gene X environment interactions (GxE), additive gene X additive gene interactions (GxG) and additive gene X additive gene X environment interactions (GxGxE). However, as Technow et al. 2021 have shown, genetic variance in elite germplasm pools is often largely statistically additive despite complex biological networks underlying the trait. Thus, the model used here will likely capture genetic effects also for more biologically complex traits in an elite germplasm context. Similarly, the existence of race-specific resistance to qualitative disease resistance sources may complicate the estimation of marker effects. There is well-documented *E. turcicum* race-specific resistance to *Ht* genes in the U.S. and Canada (Jindal et al. 2019; Weems and Bradley 2018). To avoid race-specific resistance as a confounding factor in commercial breeding field trials, disease resistance is assessed in inoculated field trials rather than under natural disease conditions. However, under natural disease pressure, the diversity of pathogen races would complicate marker effect estimation, and the model may require an additional GxE term to account for race specificity across environments.

WGOPR provides a new approach to disease resistance characterization in hybrid maize breeding by leveraging available genomic data of inbred parents in statistical methods appropriate for ordinal data. By deconstructing field-observed phenotypes into marker effects and reconstructing genome-informed disease scores, this methodology produces GECPs that retain the familiar scale used in field trials with a probabilistic interpretation. The statistical rigor and practicality of the approach ensures that breeding decisions are data-driven and disease scores are interpretable to all stakeholders.

## Notes

### Competing Interest Statement

The authors have declared no competing interest.

